# MemBright: a Family of Fluorescent Membrane Probes for Advanced Cellular Imaging and Neuroscience

**DOI:** 10.1101/380451

**Authors:** Mayeul Collot, Pichandi Ashokkumar, Halina Anton, Emmanuel Boutant, Orestis Faklaris, Thierry Galli, Yves Mély, Lydia Danglot, Andrey S. Klymchenko

**Affiliations:** Laboratoire de Biophotonique et Pathologies, UMR 7021 CNRS, Université de Strasbourg, Faculté de Pharmacie, 74, Route du Rhin, 67401 ILLKIRCH Cedex, France; University Paris Diderot, Sorbonne Paris Cité, Institut Jacques Monod, CNRS UMR 7592, 75013 Paris, France; INSERM U894, Institute of Psychiatry and Neuroscience of Paris, Membrane Traffic in Healthy and Diseased Brain, 102 rue de la Santé, 75 014 PARIS

## Abstract

The proper staining of the plasma membrane (PM) is critical in bioimaging as it delimits the cell. Herein, we developed MemBright: a family of six cyanine-based fluorescent turn-on PM probes that emit from orange to near-infrared when reaching the PM, and enable homogeneous and selective PM staining with excellent contrast in mono and two-photon microscopy. These probes are compatible with long-term live cell imaging and immunostaining. Moreover, MemBright label neurons in a brighter manner than surrounding cells allowing identification of neurons in acute brain tissue section and neuromuscular-junctions without any use of transfection or transgenic animals. At last, MemBright were used in super-resolution imaging to unravel the dendritic spines’ neck. 3D multicolor dSTORM in combination with immunostaining revealed en-passant synapse displaying endogenous glutamate receptors clustered at the axonal-dendritic contact site. MemBright probes thus constitute a universal toolkit for cell biology and neuroscience biomembrane imaging with a variety of microscopy techniques.

## Introduction

Plasma membrane (PM), in addition to its basic function of cell barrier, is a key player in crucial biological processes including cellular uptake, neural communication, muscle contraction, and cell trafficking and signalling.^1^ In the field of bioimaging, visualizing the PM is of prior importance as it delimits the cell surface. In addition, the shape of the PM directly provides information regarding the cell status such as cell division or cell death processes.^2^ Visualizing plasma membrane is particularly important in neuroscience, because it enables visualization neuronal organization and membrane trafficking involved in the synaptic transmission.^3^ Recent years have seen a tremendous expansion of fluorescence imaging techniques and tools for cellular research. In addition to genetically encoded fluorescent proteins, a number of molecular probes for monitoring cellular events have been developed.^4, 5^ The key challenge in cellular imaging is to stain cell compartments with high specificity and persistence. Although numerous efficient molecular probes have been designed to selectively stain specific cellular structures including mitochondria,^6^ lipid droplets^7^, endoplasmic reticulum^8^, nucleus^7^ and lysosomes^10–11^,there is still a demand for efficient and bright fluorescent PM markers.^12^ Fluorescently labelled lectins notably wheat germ agglutinin (WGA) and concanavalin A^9^ are popular fluorescent membrane probes. Despite their efficiency and ease of use, these probes are expensive and much larger than molecular probes. The small size of the latter, comparable to lipids, allows their precise location in the lipid bilayer, which is of key importance for studies of the lateral lipid organization of biomembranes (lipid rafts)^12, 13^ FRET with membrane proteins^14^ and superresolution imaging.^15–16^ Therefore, molecular probes are an interesting alternative to protein-based membrane markers. Although highly hydrophobic fluorophores are efficient markers of membrane models such as liposomes, they generally fail in staining the cell PM of live cells as they tend to precipitate before reaching the membrane and to quickly cross the PM to stain inner membranes.^12, 17^ Recent efforts have been made to develop specific and efficient PM probes for cell imaging with various fluorophores including chromone,^18–19^ Nile Red,^20^ styryl pyridinium,^21^ BODIPY,^22–23^ perylene,^24^ oligothiophene,^25–26^ conjugated polymers,^27^ or FITC-labelled chitosan.^28–29^ From the rapid development of multicolor imaging arose a demand for red-shifted fluorescent markers. Red-shifted fluorophores, especially near-infrared ones are highly popular for bio-imaging applications due to their long wavelength (650-900 nm) excitation and emission that reduce photodamage of biological samples, allow deep tissue penetration and avoid background signal from cell auto-fluorescence.^30–31^ We recently proposed an efficient far-red emitting PM fluorescent marker called dSQ12S based on a squaraine dye with two zwitterionic amphiphilic anchors, exhibiting superior staining compared to commonly used hydrophobic cyanines, such as DiD, PKH family, etc.^32^ Application of this unique design concept to cyanine family, highly popular in bioimaging applications,^33^ could open numerous opportunities. First, cyanines are among the brightest dyes that can be obtained in any desired emission color spanning from the visible (orange) to the near-infrared. Second, due to their capacity to form non-fluorescent *H*-aggregates, they can be transformed into fluorogenic probes.^34^ Third, cyanine dyes have been established as robust tool for superresolution microscopy (SRM),^35^ which could enable us to propose new membrane probes particularly adapted to SRM. Herein, we thus designed new membrane probes based on zwiterionic anchors and various cyanine fluorophores (Cy3/Cy3.5/Cy5/Cy5.5/Cy7/Cy7.5) using click chemistry. These probes provide very selective and bright plasma membrane staining and surpass all tested commercial dyes. They constitute a versatile tool as they allow both live, fixed, and permeabilized cells/tissue imaging using mono- or two-photon imaging. As a proof of concept we unravelled for the first time the nanoscale organisation of the axonal en-passant synapse coiling around the dendrite using 3D multicolor SRM.

## Results

### Synthesis of MemBright and spectroscopic studies

Spectroscopic properties of cyanines depend on the length of their polymethine chain, the longer chain, the higher wavelength it emits. A finer color tuning can be achieved by replacing the indole moieties by benz[e]indole ones. Herein, to obtain probes of different color, we first synthesized six di-alkyne cyanines: 3, 3.5, 5, 5.5, 7 and 7.5 (Fig. 1A). The latter were reacted with a Clickable Amphiphilic Zwitterion (CAZ) via CuAAC to provide the MemBright probes family (Fig. 1A).

**Figure 1.**
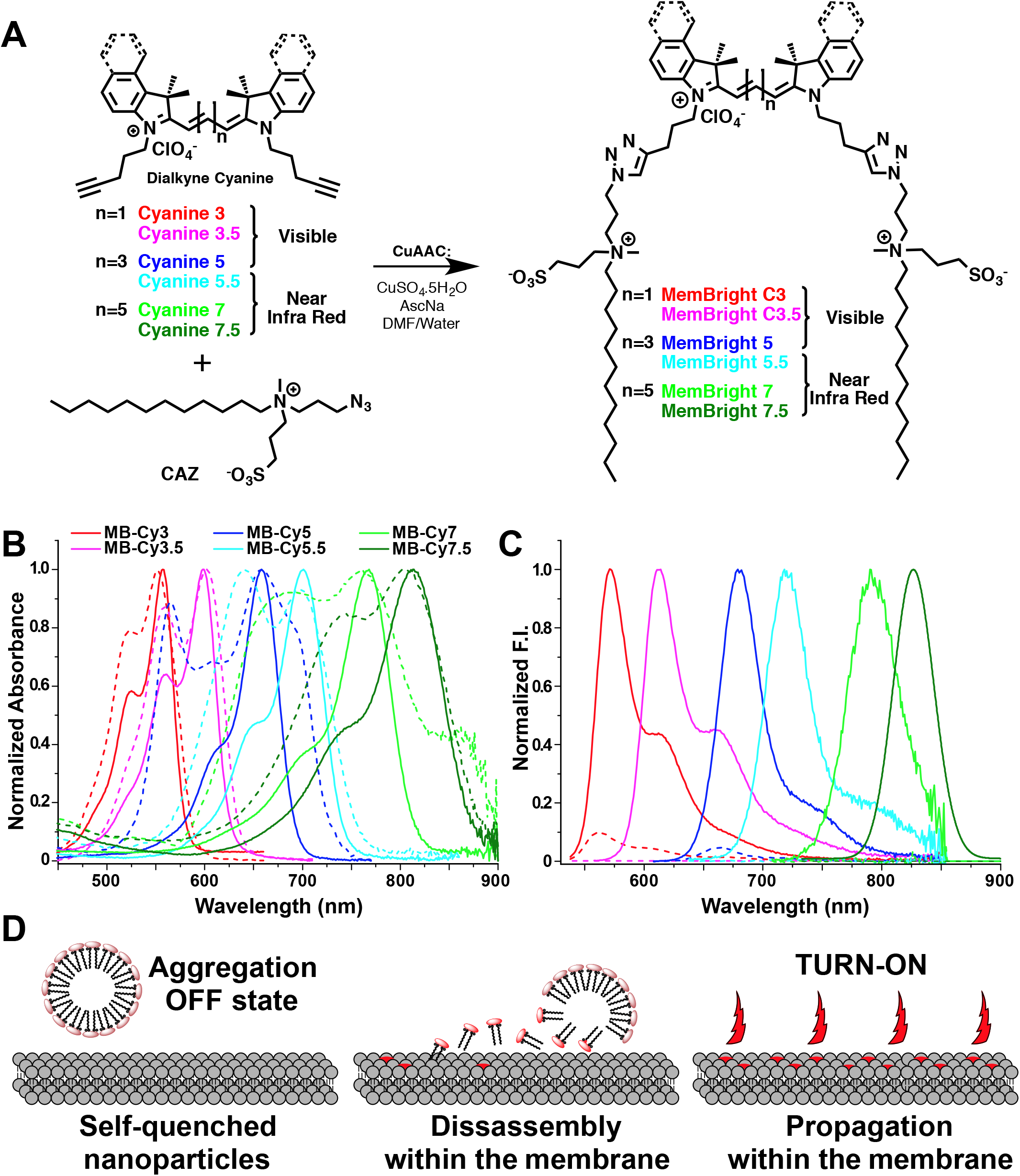
MemBright probes and their properties. Synthesis of the MemBright markers (A). Absorption (B) and emission spectra (C) of MemBright probes (200 nM) in the absence (dashed lines) or in the presence of DOPC vesicles (200 μM lipids, solid lines) in phosphate buffer 20 mM, pH 7.4. The full-corrected emission spectra beyond 850 nm were obtained by using the GaussAmp function fit. (D) Turn-on mechanism of the MemBright probes.

Spectroscopic studies of MemBright were performed in various organic solvents and buffer, as well as in the presence of Large Unilamelar Vesicles (LUVs) constituted of 1,2-Dioleoyl-sn-glycero-3-phosphocholine (DOPC) with and without cholesterol, taken as membrane models. In organic solvents, MemBright probes behaved in the same manner as their parent cyanine fluorophores. Indeed, their absorption spectra were narrow and close to the mirror image of their emission spectra, indicating no sign of aggregation (Fig. S1 to S6). Moreover, they displayed good quantum yields (0.12 – 0.66 in DMSO).

Due to their amphiphilic nature, MemBright markers are expected to form soluble aggregates in aqueous media. Indeed, DLS measurements in water showed formation of nanoparticles of 22 to 32 nm (Fig. S7). Moreover, their absorption spectra in phosphate buffer displayed a strong short-wavelength shoulder suggesting formation of non-fluorescent *H*-type aggregates, known for their blue shifted absorption^36^ (Fig. 1B). As expected, MB-Cy3 to MB-Cy5 probes displayed much lower quantum yields in buffer than in organic solvents, while MB-Cy5.5, MB-Cy7 and MB-Cy7.5 were nearly non-fluorescent in water (Table S1). In order to model the PM, we used large unilamellar vesicles (LUVs) composed of either DOPC or DOPC/cholesterol mixture (1/0.9 mol. ratio). In the presence of these LUVs, MemBright probes displayed typical absorption and fluorescence spectra of solubilized cyanine dyes (Fig. 1B and C) with quantum yield generally higher than those obtained in methanol (Table S1). The fluorescence enhancement between the aqueous media and the DOPC vesicles was 7-, 42- and 18-fold for MB-Cy3, MB-Cy3.5 and MB-Cy5, respectively, whereas for MB-Cy5.5 to MB-Cy7.5 it was much larger due to their strong quenching in aqueous media. Thus, MemBright being aggregated in water disassemble in lipid membranes (Fig. 1D), providing fluorogenic response. This mechanism is in line with the earlier data obtained with membrane probes based on Nile red^20^ and squaraine.^32^ In order to determine how fast the aggregates of MemBright disassemble in the presence of LUVs, probes were added to an excess of LUVs and their fluorescence emission intensity was followed over the time (Fig. S8). Whereas MB-Cy3 and MB-Cy5 rapidly reached their fluorescence intensity plateau in the presence of DOPC vesicles (<3 min), the other probes disassembled in a slower manner. Overall, the disassembly time increased with the polymethine chain length (Cy3 < Cy5 < Cy7) and addition of fused benzene rings (Cy3 < Cy3.5 and Cy5 < Cy5.5). We expect that more hydrophobic MemBright probes tend to form more stable aggregates, which require more time to disassemble in the presence of membranes.

### Cellular imaging

The ability of MemBright probes to selectively stain the PM was first investigated on live HeLa and KB cells. HeLa cells were chosen as they are widely used human cell lines. KB human cells (derived from HeLa cells) were chosen for their simple shape and appropriate height that facilitates PM imaging. A solution of MemBright probes (20 nM) in a serum-free medium was added to the cells and the latter were directly imaged without washing. WGA-488, a green emitting membrane marker based on a fluorescently labelled lectin (wheat germ agglutinin), was used to localize the PM. After 5 minutes, the PM displayed a clean and bright fluorescent staining with a clear co-localization of MemBright probes with WGA-488 (Fig. 2A for KB cells, Fig. S9 for HeLa cells) and high signal to background ratio (up to 18, see Table S2). Vertical projections of the confocal images confirmed perfect membrane staining by MemBright probes with no apparent internalization (Fig. 2B and S10-S11). The quality of the staining also allowed for 3D imaging of live KB cells using reconstruction of Z-stack obtained by laser scanning confocal microscopy (Fig. 2C and movies 1). These images offered high contrast and clearly revealed intercellular nanotubes, also called tunnelling nanotubes^37^ as bright thin PM structures connecting two individual cells (Fig. 2C).

**Figure 2.**
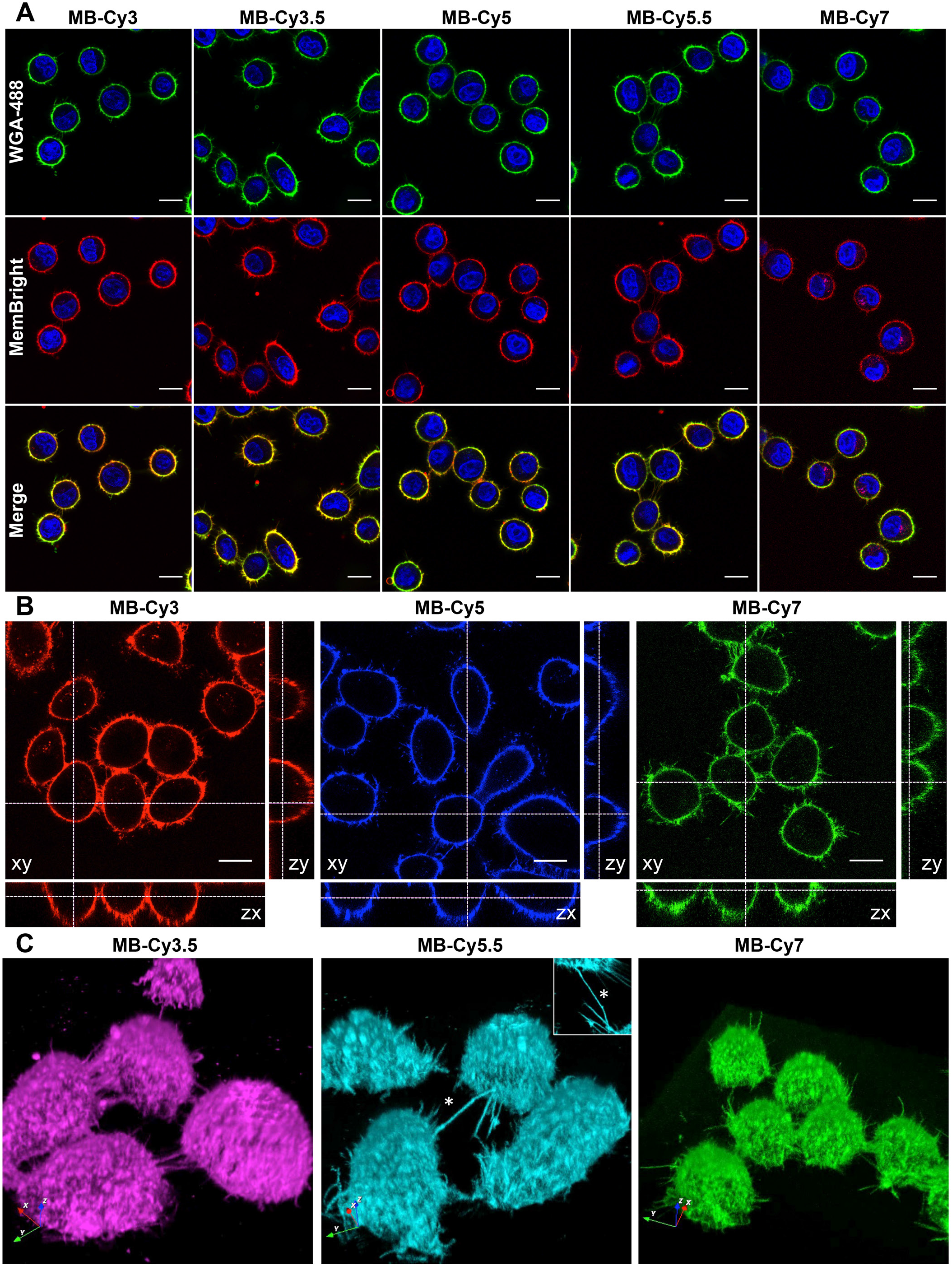
2D and 3D confocal live cell imaging using MemBright. (A) Laser scanning confocal microscopy of live KB cells labelled by 20 nM MemBright probes. Images were taken 5 min after addition of the probes (except for MB-Cy7, 15 min) without any washing step. WGA-488 was used as a co-staining marker (ex: 488 nm). Nuclear staining was done with Hoechst (5 μg/mL). (B) Orthogonal projection obtained from Z-stacks (for MB-Cy3.5 and MB-Cy5.5 see Figure S10 and S11). Scale bar is 15 μm. The microscope settings were: Hoechst 33342: ex: 405 nm with 420 and 470 nm detection range; WGA-AlexaFluor^®^488: ex: 488 nm with 500 and 550 nm detection range. MB-Cy3 and MB-Cy3.5, ex: 561 nm with 567-750 nm detection range. MB-Cy5, MB-Cy5.5 and MB-Cy7, 635 nm laser was used with 640-800 nm detection range. (C) 3D reconstruction of live KB cells stained with MemBright. Inset in B is the top view of the intercellular nanotube indicated by the white star.

According to our recent report, in similar imaging conditions, the commercial dye DID at 20 nM concentration failed to stain cellular membranes.^32^ Indeed, due to its highly hydrophobic nature, DiD is poorly soluble in aqueous media, which renders PM staining very inefficient. Another commercial probe, CellVue^®^ Claret from PKH family, is also very hydrophobic and thus, requires a specific low-salt medium for PM staining. Moreover, its cell membrane staining is heterogeneous.^32^ Therefore, commercial cyanines, such as DID (and its analogues DiI and DiO) and PKH family as well as CellMask™ membrane stains^9^ require concentration around 1-5 μM for robust PM staining. Thus, MemBright probes require ~1000 fold lower concentration compared to commercial membrane probes. Surprisingly, although MB-Cy7 also possesses a near-infrared excitation wavelength (~760 nm), it was sufficiently excited by the 635 nm laser (corresponding to 10% of its pick absorbance) and was bright enough to provide good quality 2D and 3D images.

In addition, compared to fluorescently labelled lectin WGA-488, MemBright probes displayed a more homogeneous staining of the PM when cells are confluent (Fig. S12). In the focal plane, most of the confluent cells’ PM is involved in cell-cell contacts. Within these junctions, carbohydrates located on the outer leaflet of the PM are much less accessible for the lectin than those exposed to the external medium. It was measured that in cell junctions, MemBright probes stain the PM > 3-fold more efficiently than WGA-488 (Fig. S12). This feature is due to rapid diffusion of MemBright within the whole PM, which ensures a homogeneous staining.

Next, the fluorescent staining of the PM was checked 90 minutes after addition of MemBright probes. For MB-Cy3 and MB-Cy5, the results showed that although the staining predominantly remains on the PM, internalisation partially occurred as some dots appeared in the cytoplasm, revealing the endocytosis process. In contrast, the red-shifted versions of MB-Cy3 and MB-Cy5, namely MB-Cy3.5 and MB-Cy5.5, conserved a very selective PM staining with less apparent sign of internalisation (Fig. S13). We expect that the additional fused benzene rings, which significantly increase the lipophilicity of these probes, should either favour their escape from the endocytosis process or stimulate their recycling back to the PM after endocytosis. Consequently, MB-Cy3 and MB-Cy5 will be preferred for short-term PM imaging, whereas MB-Cy3.5 and MB-Cy5.5 will be better for longer-term imaging.

At this stage of our work, it was important to evaluate the cytotoxicity of the MemBright probes. The MTT assays were performed with different MemBright probes on KB cells and the results showed no apparent cytotoxicity of the probes at concentrations up to 1 μM (Fig. S14).

Formaldehyde-based cell fixation is indispensable tool in cellular imaging. However, it leads to partial permeabilization of the PM, so that application of molecular PM probes in fixed cells is challenging. To check the possible application of the MemBright probes for imaging the PM of fixed cells, we first tried to add the dyes on fixed cells (4% PFA). Although specific staining was obtained (Fig. S15), it was noticed that prolonged time of imaging led to diffusion of the dyes within the fixed cells. However, when images obtained with WGA-488 and MemBright on fixed cells were compared, one could notice that WGA tended to stain the perinuclear envelope as well as the inner vesicles in addition to PM, whereas MemBright probes stained more selectively the PM (Fig. S16). As an alternative attempt, the dyes were added to the cells prior to fixation. This approach provided much cleaner and longer staining of the PM with only slight amount of internalized dye, demonstrating the compatibility of these probes with fixed cells (Fig. S17).

### Two-photon excitation microscopy

Two-photon excitation (TPE) microscopy imaging has received considerable attention as an advanced optical technique in the field of bio-imaging for several reasons.^38, 39^ First, TPE microscopy inherently provides three-dimensional sectioning since the excitation occurs only at the focal point of the sample. In addition, due to its localized excitation with a NIR laser, it reduces the photobleaching, the phototoxicity, and the cell auto-fluorescence. It also ensures deeper tissue penetration, which enables experiments on thicker live samples. For these reasons, there is a high demand for bright fluorescent probes with high TPE cross section values. On-going efforts have been made to design such fluorophores.^40^ Although cyanine fluorophores are widely used for bio-labelling and bio-imaging, only few studies reported on their efficiency for TPE imaging,^41, 42, 43^ giving impression that they are not well adapted for this microscopy technique. Since MemBright markers are localized in PMs in cell experiments, their two-photon fluorescence excitation spectra were measured in an aprotic solvent: DMSO (Fig. 3 A-D, top). The two-photon nature of the absorption process was verified from the quadratic dependence of the observed emission intensity *vs* excitation power (Fig. S18). To our surprise, MemBright probes displayed high two-photon action cross-section in this solvent with values reaching up to 1200 Goeppert-Mayer (GM) for MB-Cy5. TPE spectra and cross section of MB-Cy3 were found to be similar to those for the previously reported Cy3 dye.^54^ For all the MemBright probes, the maxima of TPE bands were blue shifted with respect to the one-photon absorption bands at twice the wavelength, as reported for cyanine dyes.^52^ It is remarkable that fused benzene rings significantly increased TPE cross-section for MB-Cy3.5 and shifted significantly the TPE absorption to the red for both MB-Cy3.5 and MB-Cy5.5 compared to their parent analogues (Fig. 3 A-D). In order to confirm the results obtained for the MemBright probes, TPE spectrum of the parent dye, DID (lipophilic cyanine-5), was measured in DMSO and was found to be almost identical to that of MB-Cy5 (Fig. S19). In the light of these unexpected results, MemBright probes were used in two-photon imaging experiments. Live KB cells were stained with MemBright probes without washing and imaged with two-photon excitation, chosen based on their TPE excitation spectra. High signal to noise ratio images were obtained for each of the MemBright probes with a highly selective staining of the PM (Fig. 3 A-D, bottom).

**Figure 3.**
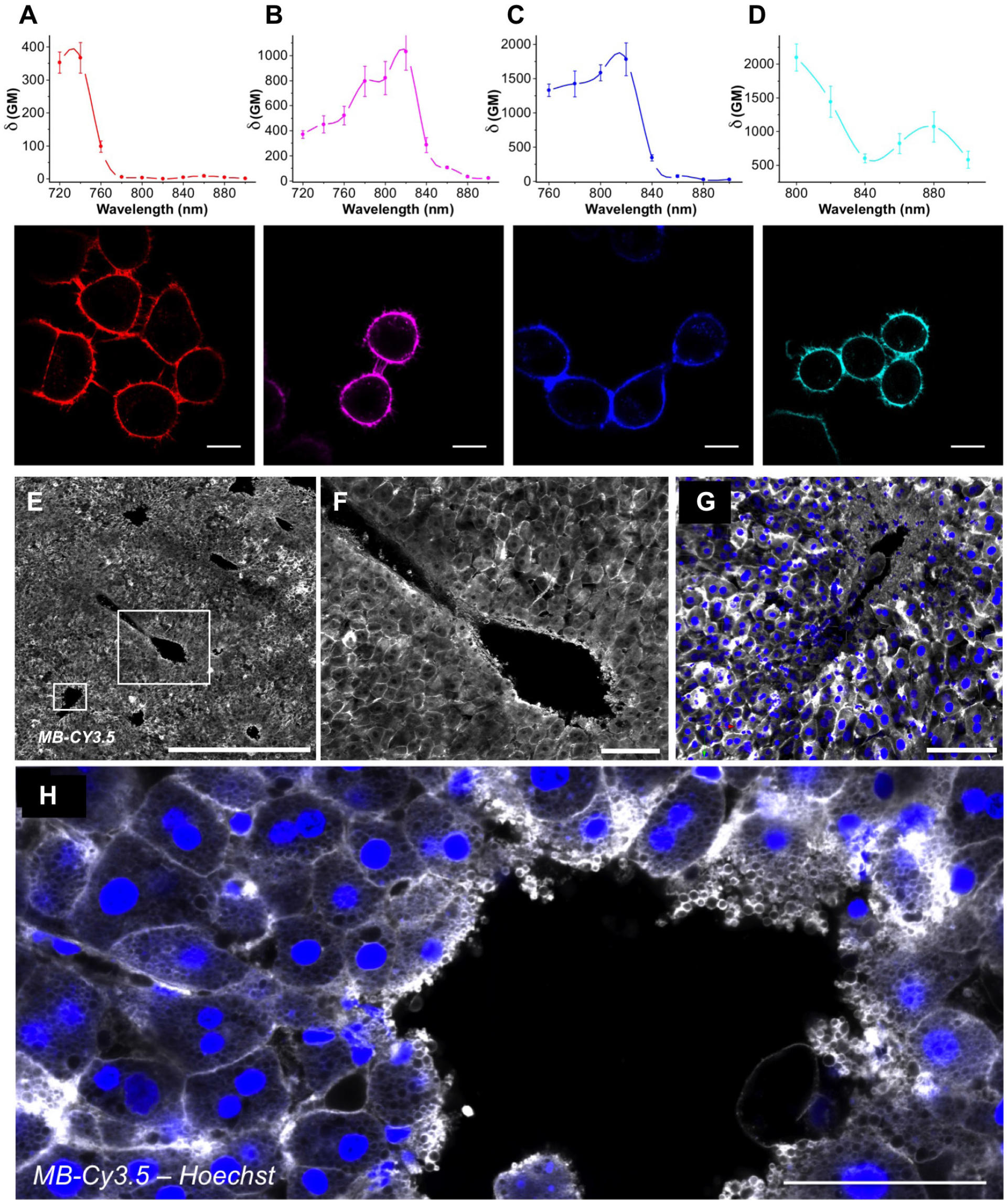
Two-photon properties and imaging. (A-D) TPE cross-section spectra (DMSO) and two-photon microscopy images of KB cells using MB-Cy3 (A), MB-Cy3.5 (B), MB-Cy5 (C) and MB-Cy5.5 (D). The images were obtained without any washing step. Conditions: MB-Cy3 at 40 nM, λEx= 760 nm (Red), MB-Cy3.5 at 40 nM, λEx= 810 nm (Magenta), MB-Cy5 at 40 nM, λEx= 810 nm (Blue), MB-Cy5.5 at 120 nM, λEx= 860 nm (Cyan). The signal was collected using a short-pass filter with 700 nm cut-off wavelength, which explained the use of MB-Cy5.5 at higher concentration. Laser power was set to 5 mW. Each image is the average of five acquisitions. Scale bar is 10 μm. (E) Tile confocal imaging of 1 mm thick liver slices incubated with MB-Cy3.5. The mosaic is composed of 4×4 fields of 20x pictures. Scale bar is 1 mm. The big and small white square correspond to the magnification in F and G-H, which shows the central vein of the hepatic lobe (G) 3D rendering of the 20x stack (101 Z slices, 55 μm thick), scale bar is 100 μm. MB-Cy3.5 is resistant to continuous illumination and allows detection of cells and tissue arrangement without any other counterstaining. (H) Confocal 63X zoom around the central vein. MB-Cy3.5 labels the PM but also the intracellular hepatic vesicles of the cells which have been cut during slicing. Scale bar is 50 μm.

### Tissue imaging

Fluorescence tissue imaging with molecular probes is more challenging as the environment is more complex and presents heterogeneity of components (cells, extracellular matrix, ducts, vessels) and fluids (e.g. plasma, blood, bile).^44, 45^ Here, applicability of MemBright probes to *ex vivo* tissue imaging was tested. As a first example, liver tissue was chosen. For this purpose, 1 mm thick blocks of rat liver were incubated for 3 h in the presence of MB-Cy5. The liver tissues were imaged by two-photon microscopy providing stacks of 50 images up to 50 μm depth. In the case of MB-Cy5 (Fig. S20) a clear PM staining was obtained and hepatocytes appeared as polygonal cells (Fig. S21). At 20 μm depth hepatocytes are well delimitated by an intense PM staining and projections on the yz and xz plans (Fig. S20B-C) showed that the PM can be stained up to 3 layers of cells in the tissue (See movie 2). The lower intensity at higher depths is clearly related to the limited diffusion of the probe through the tissue, which in principle could be improved by optimizing the staining conditions. In sharp contrast, WGA-Alexa647, used in the same conditions, showed unclear staining that appeared mainly in the nuclei (Fig. S20D-F and movie 3). Thus, MemBright is clearly advantageous compared to the commercial agent for the PM staining in tissue imaging. MemBright Cy3.5 was also very efficient in PM labelling either using two-photon excitation microscopy or conventional confocal microscopy. We tested the photo-resistance of MemBright by performing either mosaic (Fig. 3E and F) or 3D pictures (Fig. 3G and movie 4, 3D reconstruction) in large 1 mm thick liver slices. MB-Cy3.5 was resistant to continuous illumination and mosaic imaging even when acquiring 3D tile pictures reconstruction (> 100 z slices), indicating a good resistance to photobleaching. At high magnification the high signal to noise ratio of MB-Cy3.5 allowed to clearly discriminate the PM of hepatocytes and even their intracellular lipoprotein vesicles (Fig. 3H).

Encouraged by these results, we investigated on the ability of MemBright to stain the PM in brain slices. The study of neuronal arborisation has been investigated for years in various type of mutant animals using either DiI labelling thanks to biollistic approaches^46^ (DiI impregnated gold beads projected over neurons through genegun) or with the use of transgenic mice reporter (Thy-1-GFP)^47, 48^ expressing cytosolic fluorescent protein in neuronal cytoplasm. Although those technics are robust and accurate, they provide heterogeneous staining. Moreover, they are technically difficult to set up and time consuming. Herein, brain slices were incubated in the presence of MemBright before being imaged by different fluorescence imaging techniques. Taking advantage of its high two-photon absorption cross-section and red shifted emission, MB-Cy3.5 succeeded in providing good quality images of pyramidal neurons in the hippocampus of mouse brain (inset B and C Fig. 4) by two-photon excitation imaging of brain slices (Fig. 4A). The images clearly revealed the *stratum pyramidale* region (light blue layer in 5B and 5C), where the soma of the pyramidal neurons were highly stained as well as their dendrites, as thinner structures, localized in the *stratum radiatum.* 3D rendering over Z-stack (Fig. 4D) allowed visualization of the dendrite deep in the tissue as well as dendritic spines (arrows). Thus, MemBright enable fast visualization of neuronal cell layer packing as well as dendrites density.

**Figure 4.**
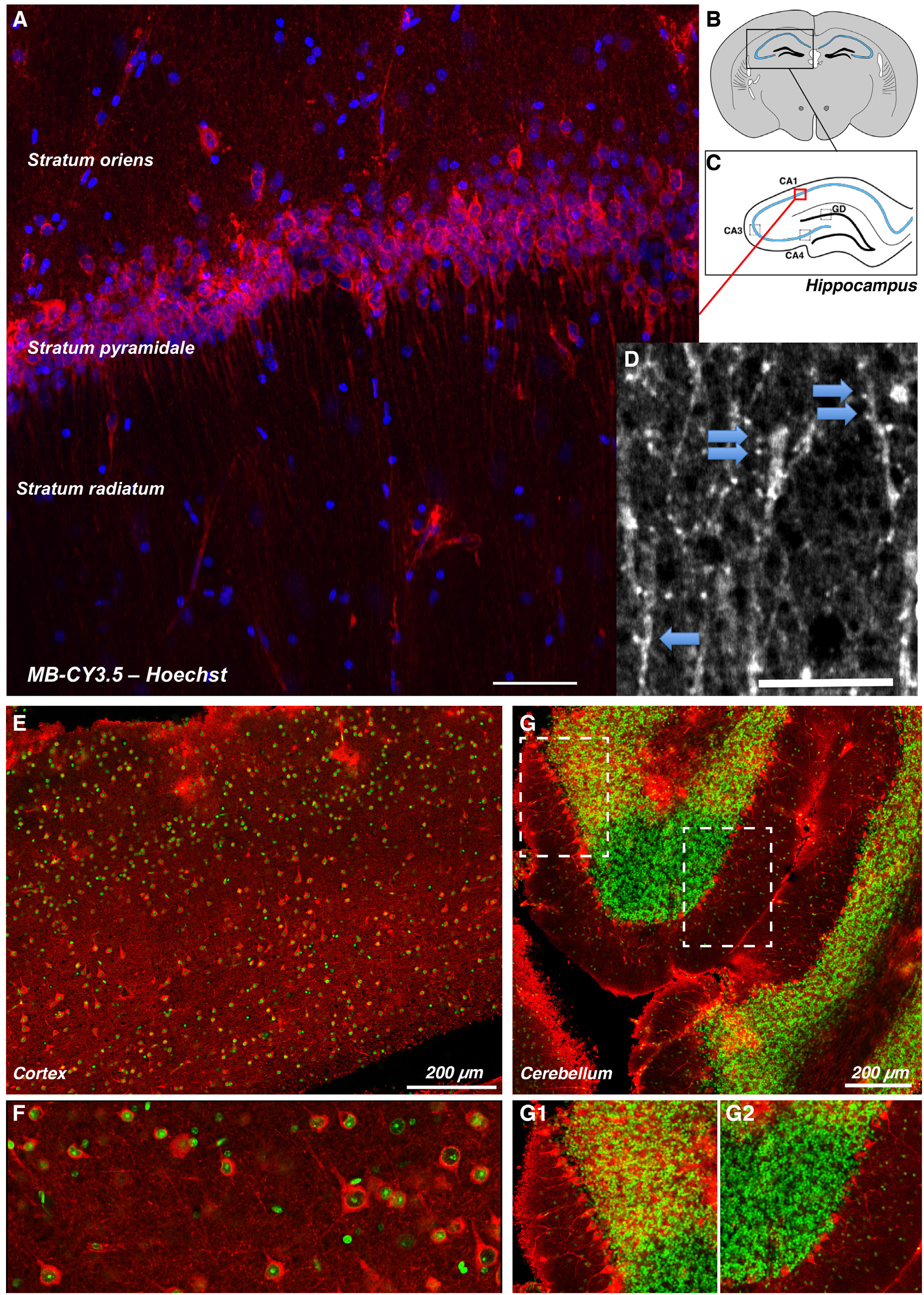
Neuronal imaging in brain slices. (A) Z projection of two-photon imaging of live 1 mm thick brain slices incubated with MB-Cy3.5 and Hoechst. 90 images were acquired every 0.4 μm and were stacked providing a 36 μm depth. Scale bar 60 μm. The red box shown in C corresponds to a zoom in stratum pyramidale of CA1 region (C) of the hippocampus (B). Two-photon imaging allows the detection of some dendritic spine in stratum radiatum (D, arrows). (E) Tile confocal imaging of 1 mm thick live cortex incubated with MB-Cy3.5. Scale bar is 200 μm. (F) Two-photon imaging showing strong labelling in cortical neurons, scale bar is 100 μm. (G) Confocal imaging of the cerebellum. G1 and G2, which shows the labelling of Purkinje cells, correspond to magnifications of the white squares in G. Scale bar is 200 μm.

Morever, these results suggest that MemBright probes are able to label neuronal cells at higher intensity than other cell types. We then checked if this characteristic was specific to hippocampal pyramidal cells, or whether MemBright could be used in other regions of the brain. 1 mm cortex (Fig. 4E-F) and cerebellum (Fig. 4G) slices were efficiently labelled and imaged either on confocal (Fig 4E) or two-photon excitation imaging (Fig 4F). Cortical neurons and Purkinje cells can be clearly seen in Fig. 4F and 4G respectively. Altogether, these results indicate that MemBright has propensity to label some major neuronal cell types.

As MemBright displayed a high affinity towards neurons, we wanted to demonstrate whether it was able to reveal motor neurons in tissue imaging. To this aim, muscle fibers with nerves were extracted from *tibialis anterior* muscles of an adult mouse and were fixed and incubated with MB-Cy5. Fluorescently labelled α-Bungarotoxin, which binds to nicotinic receptors clustered at synapse, was used to label the neurosmuscular junctions and served to visualize the whole muscle fiber. As shown in Fig. S22, the whole axon and the terminal motor nerve was clearly revealed by MB-Cy5. Both live and fixed samples could be efficiently labelled without any permeabilization. The staining was resistant enough to confocal stack imaging. 3D reconstruction allowed visualizing the nerve terminal in close apposition with the motor plate (Fig. S22B-C-D).

### Neuronal & glial cell imaging

The ability of MemBright to label polarised cells organized in networks (neurons) or in confluent layers (pavement of astrocytes) was investigated. In this endeavour, we used hippocampal neuronal cells which can be seed and fed on astrocytes feeder layer.^49^ Primary hippocampal cultures are efficiently labelled with any member of the MemBright family (Fig. 5, for MB-Cy5.5 see Fig. S23), revealing neuronal cells lying over astrocytes. This experiment confirmed what was previously noticed in tissue imaging, that neurons appeared in a much brighter manner than glial cells, suggesting a preferential staining of neurons by MemBright. Although MemBright probes were used at very low concentration (typically, 20 nM directly in the imaging medium), they allow revealing even very fine protrusion including astrocytic filopodia (Fig. 5A3, A4, E, E1) or fine dendritic processes (Fig. 5A, B, C and D). Unlike the use of fluorescent proteins, this method leads to the labelling of all cells in live microscopy without any transfection steps that are poorly efficient and can be deleterious in neuronal cells.

**Figure 5.**
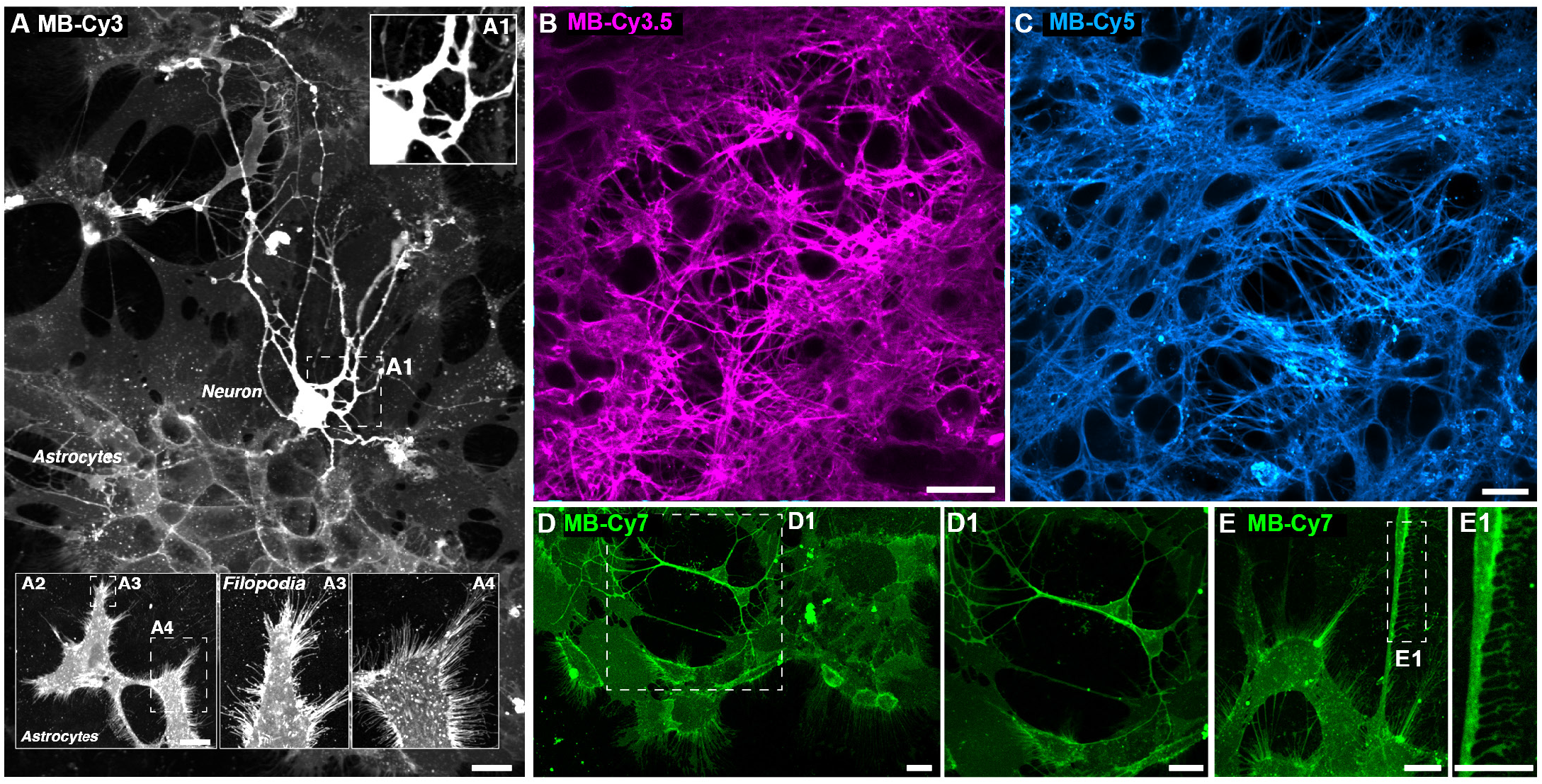
Confocal microscopy of hippocampal primary neurons and astrocytes labelled with MemBright probes. Images were taken 10-15 minutes after addition of the probes (20 nM) without any washing step. (A) Hippocampal neurons appear in a brighter manner and can be identified on astrocytes the latters being either confluent (as shown in A) or isolated (A2). Zooms in the PM show how the MemBright allow detection of very fine membrane protrusion in astrocytes (filopodia in A3 or A4) or in neuronal cells (dendritic processes in A1). (B-C) Live dense neuronal network is labelled with 20 nM MB-Cy3.5 (B) or 20 nM MB-Cy5 (C) and imaged with spinning disc microscopy. (D-E) Example of astrocytic layers and isolated neurons labelled with 200 nM MB-Cy7 excited with 633 nm laser and imaged with laser confocal scanning microscopy. Even in the near infrared region, membrane details can be unravelled with the probe both on neuronal (D1) and glial cells (see fine filopodia in E1). Scale bars are 20 μm.

Live cells could be labelled and imaged on confocal or spinning disc microscopy in combination with other vital staining (Fig. S23-S24). Very interestingly, labelled live cells could be kept with the dye during several days without any visible detrimental effects. Indeed, neuronal culture composed of hippocampal primary neurons and glial cells was labelled with MemBright and imaged over several hours under confocal illumination. As seen in supplementary movie 5, the brightness of MemBright and its low toxicity allowed to monitor the cells with only 0.2% power of 561 nm laser line over 13 h without any loss of membrane staining. Fine neuronal filopodia and widespread glial lamellipodia could be easily tracked thanks to MemBright imaging. MemBright is thus able to provide an efficient membrane labelling over long periods even at 37°C. We then compared MemBright labelling on live neurons with other classical membrane probes. As expected from HeLa and KB cells labelling, the WGA staining on neurons was very bright and homogenous from cell to cell (Fig. S25A). However when inspecting carefully the PM, we noticed dashed staining along the PM, impeding fast segmentation of cell shape (Fig. S25B). This might be attributed to the non-homogeneous expression of glycosylated protein at the PM. To circumvent this problem we reasoned that staining by lipidic dyes should be more homogeneous. Consequently, MemBright was compared to DID (Fig. 25C-F) over neuronal network. Although DiD labelling, compared to WGA, was more homogeneous along the PM and allowed perfect visualization of the cell shape, staining intensity from cell to cell was variable ranging from very faint (white arrowhead in Fig. S25C-D) to very bright (red arrow head in Fig. S25C-D). MemBright was then compared to mCLING, a polylysine-based membrane probes developed for super-resolution imaging.^50^ For this purpose, we strictly followed the protocol provided by the developers of mCLING.^51^ As shown in Fig. S26A, after 10 min incubation, mCLING labelling is mostly punctiform and seemed to be mainly internalized in intracellular vesicles as shown previously.^50^ Conversely, MemBright was still present at the PM and clearly allowed the identification of neuronal cell bodies (Fig. S26B) as well as axonal or dendritic networks.

### Monitoring membrane trafficking and cellular architecture with endogenous proteins

We then explored whether MemBright probes allowed tracking of endocytic vesicles. We thus performed incubation of MemBright at 37°C with an antibody directed against L1-CAM, an adhesion protein expressed at the cell surface and known to be recycled by trafficking vesicles.^52^ As shown in Fig. 6A1-A3, MemBright colocalized perfectly with L1-CAM at the cell surface. Live antibody uptake (Fig. 6A4-A6) showed that endocytic vesicles containing L1-CAM are also labelled with MemBright, indicating that a portion of MemBright can be internalised to monitor endocytosis. It is noteworthy that vesicles stained with MemBright are more numerous than vesicles containing L1-CAM indicating that MemBright can be used to track various endocytic pathways. Additionally, a high amount of MemBright is still present at the PM surface, which can help assessing membrane morphology and monitoring endocytosis in the same channel.

**Figure 6.**
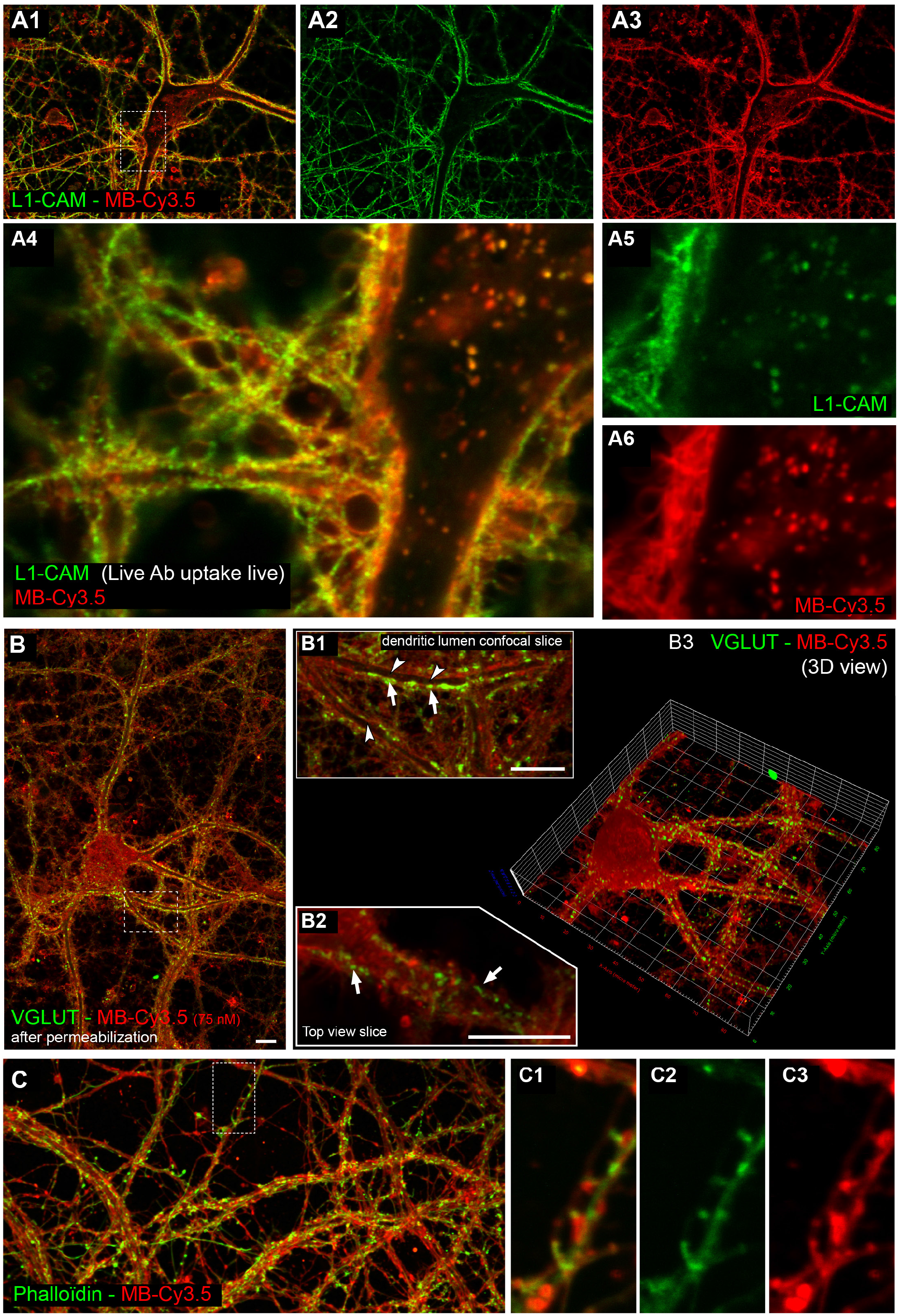
Multicolor imaging using MemBright and labelled endogenous proteins. (A) Confocal imaging of primary hippocampal neurons incubated with MB-Cy3.5 (red in A1-A3, A4 and A6) and L1-CAM mAb (Green in A1-A2, A4 and A5) for 10 min at 37°C. MB-Cy3.5 allows the identification of the PM both on the cell body and dendrites as shown with colocalization of the cell surface L1-CAM adhesion molecule in A1. Internalised vesicles can be tracked with L1-CAM antibody (zoom in A5) or with MB-Cy3.5 (A6) for a wider identification of different endocytic pathways. (B) Confocal imaging of primary hippocampal neurons incubated with MB-Cy3.5 (B1-B3) for 10 min at 37°C then fixed, permeabilised and incubated with pAb to VGLUT (Green). MB-Cy3.5 allows the identification of the PM both on confocal section passing either trough the dendrite (B1) or below the dendrite (B2) which can be visualized in 3D thanks to stack reconstruction (B3). Presynaptic boutons labelled with VGLUT (arrows) can be seen in contact on the dendritic membrane. Even the dark lumen of dendrite (arrowhead) can be visualized. (C) Confocal imaging of primary hippocampal neurons incubated with MB-Cy3.5 (red) and phalloidin A488 (green) showing that dendritic spine head/neck can be seen with both dyes.

Using a PM sensor in conjunction with classical immunochemistry would avoid time-consuming transfection of GFP reporters that are usually not efficient and very difficult in primary cells. As discussed previously, fixation followed by permeabilization leads to leakage of the dye out of the PM. Here, after optimizing the permeabilization process we used MemBright labelling in combination with primary VGLUT antibody to unravel the glutamatergic synaptic vesicles within axonal presynaptic sites. As shown in Fig. 6B, we succeeded in maintaining MemBright labelling after fixation, allowing the detection of VGLUT synapse (arrows in 9B1 and 9B2) contacting dendrite labelled with MemBright. Confocal section allowed the visualization of either dendritic membrane from a top external view (in 9B2) and from a crossing section revealing the dendritic lumen (9B1). 3D-view of a Z-stack led to a clear identification of the neuronal cell body and dendrites (in red) contacted by several glutamatergic synapses (in green).

Dendritic spines are known to be very dynamic tiny structure that change shape during memory,^53^ therefore we decided to tackle this detection with MemBright. The identification of the spine shape is usually done with phalloidine on fixed samples or with transgenic fluorescent mice or transfection on live neurons. Double labelling with phalloidine and MemBright on live samples showed that both dendritic spine head and neck can be efficiently labelled by MemBright with a preserved labelling after fixation (Fig. 6C1-A3).

### Unravelling synapse and dendritic spine morphology using super-resolution imaging

Over the last decade, super-resolution fluorescence microscopy has pushed the diffraction limit in cell imaging.^54^ Among the available techniques, Stochastic Optical Reconstruction Microscopy (STORM)^55^ and PhotoActivation Localization Microscopy (PALM)^56–57^ are based on successive activation, imaging and high precision localization of photoswitchable or blinking fluorophores. Here, MemBright probes were used at 20 nM to stain fixed HeLa cells which were imaged by Total Internal Reflection Fluorescence (TIRF) Microscopy and high-resolution dSTORM. Among the developed probes, MB-Cy3.5 was selected as the most efficient for dSTORM imaging. After an initial bleaching phase, MB-Cy3.5 exhibited a stable blinking. A series of 6000-10000 frames were acquired and high-resolution images were reconstructed (Fig. S28). The localization precision depending on the intensity of the background signal ranged between 17-20 nm (~700 photons detected). dSTORM image of MB-Cy3.5 labelled PM clearly showed an increase of the resolution compared to the standard TIRF image (Fig. S27A-B). Indeed, the intensity plot profile of a cell filopodium evidenced a decrease of the thickness from 430 nm to 120 nm that corresponds to a 3.5-fold increase of the image resolution (Fig. S27C).

With this tool in hand, we then assessed if MBCy3.5 could reveal dendritic spine morphology on fixed samples in 3D STORM microscopy. First, we showed that MemBright efficiently labelled dendritic spine neck and head, allowing the identification of mushroom spines at either 9 (Fig. 7A-C, and movie 6) or 14 days *in vitro* (Fig. 7D). 3D imaging with color-coded depth allowed the visualization over several microns from back (Fig. 7B) to front view (Fig. 7B and 10D). As a comparison, we assessed the ability of our previously developed PM probe dSQ12S to provide dSTROM imaging in similar conditions. Although dSQ12S successfully stained the PM of neurons in widefield microscopy, it rapidly photobleached (Fig. S28) and did not blink sufficiently to provide dSTORM super resolution images (Fig. S29). Then, MemBright was used as a reference PM probe in multi-color STORM microscopy after fixation and permeabilization. As shown in Fig. 7E-G, MemBright allowed visualizing both axonal (Fig. 7E-G-I) and dendritic processes (Fig. 7E-F-H). Since MemBright stains axonal processes in a much brighter manner than dendrite, it first allowed us to visualize axon coiling around a dendrite in the first thousands of pictures; then, longer acquisition revealed the whole dendritic processes. Additional immunostaining of AMPA glutamate receptors (GluR2 subunit) revealed that glutamate receptors were accumulated at the axon-dendritic junctions. Altogether, these experiments showed that MemBright can be used as reference membrane markers in both conventional and STORM microscopy on either live or fixed samples.

**Figure 7.**
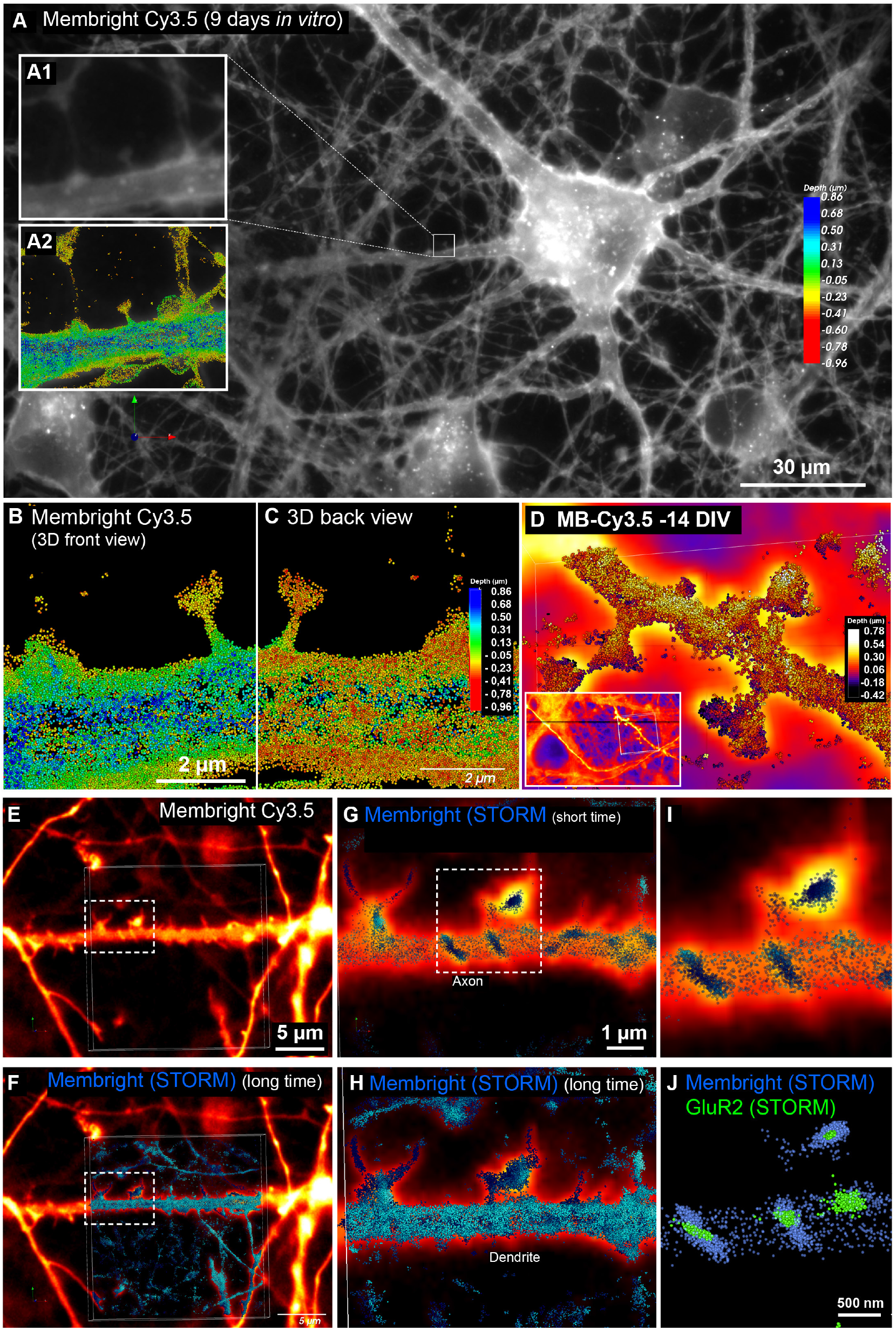
STORM microscopy of neurons using Membright Cy3.5. Imaging of hippocampal neurons labelled with MemBright after 9 (A-B-C) or 14 (D) days in vitro. Spines can be identified in widefield microscopy (A1) and then super-resolved using STORM (A2-B-C). 3D STORM microscopy allowed visualizing the spine from the front (B) or the back view (C). See supplementary movie 6 for animated views. MemBright Cy3.5 allows the visualization of the PM through several microns thanks to biplane module, depth is color-coded both in B-C-D (rainbow LUT). Dendritic and axonal profiles can be visualized in widefield microscopy (in E). Axonal processes coil around the dendrite (dashed box in E, zoom in G and I) and can be reconstructed in the first 3D-STORM images due to intense blinking. After several hundred pictures, the dendritic shape can be reconstructed in F (zoom in H). Thanks to permeabilzation resistance, MemBright (blue) can be used in dual color immunofluorescence to visualize glutamate AMPA receptors clusters (green) aggregated at the dendrite in front of axonal profile (J) with interleaved 3D STORM stack.

## Discussion

In this work, we developed a family of bright PM probes based on cyanines, so-called MemBright, of six different colors and demonstrated its versatility for imaging the PM by several imaging techniques. Taking advantage of the symmetric structure of cyanines, they were modified with two amphiphilic zwiterionic anchors ensuring the selective and persistent staining of the PM. In aqueous media, these probes form non-fluorescent dark aggregates, whereas in the presence of lipid membranes, these aggregates dissociate into highly fluorescent molecular species solubilised in the bilayer. This turn-on capability of MemBright probes enabled cellular imaging without any washing step with high signal to noise ratio in both live and fixed cells. A structure-property relationship was established as the most lipophilic probes (MB-Cy3.5, MB-Cy5.5 and MB-Cy7.5) tend to label the PM in a slower manner (15 minutes vs 5 minutes) but, in the meantime, provide longer-term staining of the PM. Owing to the high brightness of their cyanine fluorophore and efficient membrane staining, MemBright probes can be used at concentrations as small as 1 nM, which is >1000-fold lower than those commonly used for long-chain cyanines, like DID, PKH family or mCLING. To get sufficient signal other probes are used at higher concentrations but can then become toxic for cells as shown for mCLING above 2 μM.^51^ Moreover, in confluent cells, MemBright probes were found to stain the PM in a more homogeneous manner than the fluorescently labelled WGA, a protein-based membrane marker, because the former are small molecules that can diffuse rapidly within the lipid membrane and unlike WGA are not dependent on the heterogeneous expression of glycan at the cell surface. This property of MemBright probes can be a real asset for imaging PM in 3D cell cultures, tissues and small animals, featuring tight packing of the cells. MemBright probes were also found to be efficient in two-photon excitation imaging, with two-photon cross-section reaching 1800 GM for Cy5-based probe, and served to image deep PM in liver and brain tissues. In the field of neuroimaging, MemBright probes were successfully applied for imaging live and fixed neuronal networks, showing excellent imaging contrast of the PMs of the primary neurons together with astrocytes. This is a really powerful advantage over transfection method, which are usually deleterious for the cells and which allow the detection of only 5-10 % of the neurons within a population. This has prompted the user in the past to use transgenic reporter mice, such as THY1-fluorescent transgenic lines.^58^ However, this method is almost reserved to the mouse species since transgenic rat, marmoset or other mammal animal models are still rare. The MemBright are here usable whatever the species and eventually on human samples coming from chirurgical resections. Moreover, due to the wide range of available colors, MemBright can also be used in combination with green or red transgenic mice using complementary red, deep-red or near infrared MemBright. Other alternatives used to label neurons in brain tissue are biocytin injection^59^ and DiI, DiO and DiD tissue-labelling pastes or crystals.^60^ Whereas the injection needs a fluorescent streptavidin application to reveal the neurons, the second one rely on insoluble dyes in water which can be problematic for application.^61^ Indeed those products are usually placed via a needle into the brain area or by using gene gun. This usually results in a non-homogenous dispersion of the dye leading to a very bright local fluorescence. MemBright are water-dispersible and thus can be used either incubated on cultured cells or directly on slices, which does not require any special skills. Moreover they can be visualized directly without any additional revealing steps thanks to their intrinsic fluorescence. At last MemBright probes combine the capability of the Neurotrace™ dye^9^ for neuronal soma and Fluoromyeline™ dye^9^ for labelling myelin in spinal cord.^62^ Indeed, MemBright stains well the cell soma as well as the dendrites (Fig. 5 and 10), axons (Fig. 7 and 10) and commissural and association fibers (*corpus callosum*, *fimbria* – data not shown). Very interestingly, it appeared that MemBright markers preferentially stain the PM of neurons with a higher intensity compared to surrounding cells present in the brain (e.g. glial cells). This tendency allowed imaging neurons in various regions of the brain tissue with high signal to background ratio. Additionally, MemBright enables high-contrast imaging of motor neurons *in vivo,* which opens perspectives in the important field of nerve-specific fluorescence guided surgery.^63^

Finally, we showed that MemBright Cy3.5, owing to its capacity to blink, enables STORM-type super-resolution microscopy. To our knowledge, MemBright are the first plasma membrane probes operating at nanomolar concentrations in live or fixed samples using conventional confocal, two-photon as well as super resolution (STORM) microscopy. Since MemBright is compatible with cell fixation and permeabilization, it can be used in combination with immunofluorescence and open a wide avenue to multicolor imaging at the nanoscopic resolution. Several teams are currently trying to decipher the nanoarchitecture of molecular complexes with STORM microscopy.^64 65 66 67 68^ It is thus crucial to get a plasma membrane staining to precisely localize these proteins in their plasma membrane environment in order to map their cellular distribution. Additionally, this is of paramount importance in the context of 3D reconstruction. Most of actual studies lacki of membrane staining as reference and display dotted lines indicating the putative location of cell limits. We provide here the first blinking membrane probe which is alsoresistant enough to permeabilization and reducing buffer to allow such a use in multi-color STORM immunostaining. We showed for the first time the nanoscale organization of both axonal and dendritic compartments in combination with endogenous glutamate receptors clusters in multi-color 3D STORM. Glutamate receptors are clearly aggregated at contact site between axon and dendrites as well as at extrasynaptic sites. This membrane reference thus allows the fine molecular mapping of endogeneous proteins at the nanoscale level within the cellular context that was, up to now, only possible with EM microscopy. In conclusion, due to the availability of six different colors within 550-850 nm spectral range, combined to their ease of use, their brightness and compatibility with different microscopy techniques, the MemBright family appears as a powerful toolbox for bio-membrane imaging in cell biology and neuroscience.

## Material and Methods

### Abbreviations

PM: Plasma Membrane; WGA-488: Wheat Germ Agglutinin-Alexa-488; FRET: Förster Resonance Energy Transfer; FITC: Fluorescein IsoThioCyanate; CuAAC: Copper(I)-catalyzed alkyne-azide cycloaddition; LUV: Large Unilamellar Vesicles; DOPC: Dioleoylphosphatidylcholine; DMSO: Dimethyl Sulfoxyde; PB: Phosphate Buffer; Chol: Cholesterol; DLS: Dynamic Light Scattering; PFA: Paraformaldehyde; GM: Goeppert-Mayer; STORM: Stochastic Optical Reconstruction Microscopy; PALM: PhotoActivation Localization Microscopy; TIRF: Total Internal Reflection Fluorescence; RT: Room Temperature; MEA: mercaptoethanol amine.

### Synthesis of MemBright

Synthesis, protocols, characterizations and spectra are described in the supporting information. NMR spectra were recorded on a Bruker Avance III 400 MHz spectrometer. Mass spectra were obtained using an Agilent Q-TOF 6520 mass spectrometer.

### Lipid Vesicles

Dioleoylphosphatidylcholine (DOPC) and cholesterol were purchased from Sigma-Aldrich. Large unilamellar vesicles (LUVs) were obtained by the extrusion method as previously described.^69^ Briefly, a suspension of multilamellar vesicles was extruded by using a Lipex Biomembranes extruder (Vancouver, Canada). The size of the filters was first 0.2 μm (7 passages) and thereafter 0.1 μm (10 passages). This generates monodisperse LUVs with a mean diameter of 0.11 μm as measured with a Malvern Zetasizer Nano ZSP (Malvern, U.K.). LUVs were labelled by adding 5 μL of probe stock solution in dimethyl sulfoxide to 1-mL solutions of vesicles. A 20 mM phosphate buffer, pH 7.4, was used in these experiments. Molar ratios of probes to lipids were generally 1 to 500-1000.

### Spectroscopy

Absorption spectra were recorded on a Cary 4000 spectrophotometer (Varian) and fluorescence spectra on a Fluoromax 4 (Jobin Yvon, Horiba) spectrofluorometer. Fluorescence emission spectra were systematically recorded at room temperature, unless indicated. All the spectra were corrected from wavelength-dependent response of the detector. The fluorescence and absorption spectra of the corresponding blank suspension of lipid vesicles without the probe was subtracted from these spectra. Concentrations of dyes were 400 nM for MB-Cy3 and MB-Cy3.5, 280 nM for MB-Cy5 and MB-Cy5.5, 220 nM for MB-Cy7 and 600 nM for MB-Cy7.5. Quantum yields were determined by comparison with a reference according to their excitation and emission wavelengths: Rhodamine 101 in EtOH^70^ was used as the reference for MB-Cy3, Cresyl Violet in EtOH^70^ for MB-Cy3.5, DID in MeOH^71^ for MB-Cy5, Rhodamine 800 in EtOH^72^ for MB-Cy5.5 and Indocyanine green in MeOH^73^ for MB-Cy7 and MB-Cy7.5.

### DLS measurements

The sizes of the particles formed by MemBright in water were measured by Dynamic Light Scattering (DLS) using a Malvern Zetasizer Nano ZSP (Malvern, U.K.). Concentration of the dye was 5 μM. The size of particles formed by MB-Cy5 and MB-Cy5.5 could not be measured due to the wavelength of the DLS laser (633 nm).

### Cellular studies

HeLa cells (ATCC^®^ CCL-2) were grown in Dulbecco’s modified Eagle medium (DMEM, Gibco-Invitrogen), supplemented with 10% fetal bovine serum (FBS, Lonza) and 1% antibiotic solution (penicillin-streptomycin, Gibco-Invitrogen) at 37° C in humidified atmosphere containing 5% CO_2_. KB cells (ATCC^®^ CCL-17) were grown in minimum essential medium (MEM, Gibco-Invitrogen) with 10% fetal bovine serum (FBS, Lonza), 1% non-essential amino acids (Gibco-Invitrogen), 1% MEM vitamin solution (Gibco-Invitrogen), 1% L-Glutamine (Sigma Aldrich) and 0.1% antibiotic solution (gentamicin, Sigma-Aldrich) at 37° C in humidified atmosphere containing 5% CO_2_. Cells were seeded onto a chambered coverglass (IBiDi^®^) at a density of 1×10^5^ cells/well 24h before the microscopy measurement. For a nuclear staining, the medium was replaced by Hoechst 33342 (ThermoFisher, 5 μg/mL) in Opti-MEM (Gibco-Invitrogen) and the cells were incubated for 10 minutes at 37°C. For imaging, the medium was removed and the attached cells were washed with HBSS (Gibco-Invitrogen) three times. Then, a freshly prepared solution of MemBright in HBSS (typically 20 nM) was quickly added to the cells without any washing step. Prior to imaging, PM was costained by addition of wheat germ agglutinin-Alexa488, WGA-AlexaFluor^®^488 (1 mg/mL in water) at a final concentration of 5 μg/mL. Confocal microscopy experiments were performed by using a Leica TCS SPE-II with HXC PL APO 63x/1.40 OIL CS objective. The microscope settings were: 405 nm laser for excitation of Hoechst 33342, emission was collected between 420 and 470 nm; 488 nm laser for excitation of WGA-AlexaFluor^®^488, emission was collected between 500 and 550 nm. For excitation of MB-Cy3 and MB-Cy3.5, 561 nm laser was used with 567-750 nm detection range. For excitation of MB-Cy5, MB-Cy5.5 and MB-Cy7, 635 nm laser was used with 640-800 nm detection range. The images were processed with the ImageJ software. In the case of fixed cells 1) fixation followed by staining. The HeLa cells seeded onto the 1 mL Ibidi^®^ Chamber were washed once with D-PBSx1 (Lonza) and fixed in 4% paraformaldehyde (PFA - Electron Microscopy Sciences) at room temperature during 10 min. The fixed cells were washed three times with D-PBS to remove the excess of PFA before usual staining and imaging. For an efficient staining on fixed cells, 2) staining followed by fixation. The cells seeded onto the 1 mL Ibidi^®^ Chamber were stained with 20 nM of MemBright for 10-15 minutes at room temperature, the excess of MemBright was washed with HBSS and the cells were fixed with 4% paraformaldehyde in HBSS for 10 minutes at room temperature before being washed 2 times with HBSS.

### Neuromuscular junction preparation

Dissected hindlimb tibialis anterior (TA) muscles from adult C57BL6 mice were fixed (4% PFA in PBS) for 20 minutes at room temperature and further incubated overnight at 4°C with MB-CY5 (200 nM), Hoechst and alpha-bungarotoxin-Alexa594. After three washed in PBS, mounted under glass in Vectashield (Vector Labs, Burlingame, CA). Incubation of MB-Cy5 for 30 min on non-fixed preparations also gave good results.

### Cytotoxicity assay

Cytotoxicity assay of the MemBright dyes was quantified by the MTT assay (3-(4,5-dimethylthiazol-2-yl)-2,5-diphenyltetrazolium bromide). A total of 1×10^4^ KB cells/well were seeded in a 96-well plate 24 h prior to the cytotoxicity assay in Dulbecco’s Modified Eagle Medium (Gibco Lifetechnologies-DMEM) complemented with 10% fetal bovine serum, Penicilin (100 UI/mL), Streptomycin (100 μg/mL), L-Glutamine (2 mM) and were incubated in a 5% CO_2_ incubator at 37°C. After medium removal, an amount of 100 μL DMEM containing 1000 nM, 200 nM or 20 nM of MemBright (MB-Cy3, MB-Cy3.5, MB-Cy5, MB-Cy5.5 and MB-Cy7) was added on the KB cell and incubated during 1 h at 37°C (5% CO_2_). As control, for each 96-well plate, the cells were incubated with DMEM containing the same percentage of DMSO (0,5% v/v) as the solution with the tested dyes or with Triton 1% as a positive control of cytotoxicity. After 1h of dye incubation, the medium was replaced by 100 μL of a mix containing DMEM + MTT solution (diluted in PBS beforehand) and the cells were incubated during 4 h at 37°C. Then, 75 μL of the mix was replaced by 50 μL of DMSO (100%) and gently shaken for 15 min at room temperature in order to dissolve the insoluble purple formazan reduced in living cells. The absorbance at 540 nm was measured (absorbances of the dyes at 540 nm were taken into account). Each concentration of dye was tested in sextuplicate in 3 independant assays. For each concentration, we calculated the percentage of cell viability in reference of the control DMEM+ 0,5% DMSO.

### TPE measurements

Two-photon absorption cross-section measurements were performed using Rhodamine B in methanol as a calibration standard according to the method of Webb *et al.* Two-photon excitation was provided by an InSight DS + laser (Spectra Physics) with a pulse duration of 120 fs. The laser was focused with an achromatic lens (f = 2 cm) in a cuvette containing the dye (0.2–1.0 μM in DMSO) and the spectra were recorded with a fibered spectrometer (Avantes) by collecting the fluorescence emission at 90° with a 20 × Olympus objective.

### Two-photon imaging

Two-photon fluorescence microscopy imaging was performed by using a home-built two-photon laser scanning setup based on an Olympus IX70 inverted microscope with an Olympus 60x 1.2NA water immersion objective.^74^ Two-photon excitation was provided by an InSight DS + laser (Spectra Physics), and photons were detected with Avalanche Photodiodes (APD SPCM-AQR-14-FC, Perkin-Elmer) connected to a counter/timer PCI board (PCI6602, National Instrument). Imaging was carried out using two fast galvo mirrors in the descanned fluorescence collection mode. Typical acquisition time was 50 s with an excitation power of 5 mW at the laser output. The images were processed with ImageJ.

### Tissue imaging

Rat tissue: Fresh samples of rat liver tissue were cut (2×5 mm, 1 mm thickness) and kept in Krebs solution at RT. Before staining, the samples were washed 3 times with Krebs solution at RT. For the staining, the tissue samples were placed 3 h at RT in 1 mL of freshly prepared solution of dye (MB-Cy5: 5 μM in HBSS, WGA-647: 10 μg/mL in HBSS). The tissue samples were washed 3 times with HBSS before being placed in a iBiDi dish containing 3 mL of HBSS, and were then flattened by adding a glass slide on the top of it. The samples were excited at 810 nm with a power of 5 mW for both MB-Cy5 and WGA-647. 50 frames (60 × 60 mm, 512 × 512 pixels) were collected with a depth step of 1 μm, providing stacks of 50 μm depth. Each frame was scanned once with a speed of 4 ms per pixel. The images were processed with ImageJ.

Mice tissue: C57BL6/J mice were maintained on a 12 hour light-dark cycle with *ad libitum* access to food and water. All animal work was conducted following protocol approved by ethical committee. Adult C57BL6 mice were euthanized by CO_2_ administration. Liver and brain were immediately dissected and washed in PBS at 4°C, before being sliced in 1 mm slices on a Leica Vibratome. Tissue slices were placed either at RT or at 4°C in 1 mL of freshly prepared solution of dye (MB-Cy3.5, Cy5 and Cy5.5: 5 μM in PBS) for a variable period (from 1 to 24 h). The tissue samples were washed 3 times with PBS before being placed in a homemade glass chamber, allowing the imaging on both side of the slides. Briefly a 1 cm hole was breached in a glass slide thanks to a diamond drill bit. The 1 mm tissue slice was inserted in the hole, covered by a coverslip (170 nm, #1.5) on each slide, and sealed with Picodent twinsil. The slices were excited on a LavisionBiotec two-photon Microscope at 800 nm with a TiSa laser (Insight Spectra Physics (690-1300 nm) for both MB-Cy3.5 and Hoechst. Each frame was scanned twice (line average 2) with a speed of 0.4 μs per pixel using a 20x plan Apo Chroma Zeiss (NA:1) water-immersion objective. The fluorescence signal was detected with GaAsP detectors. Confocal microscopy was performed on a Leica SP5 microscope (405 nm (25 mW); Argon Krypton 488 (40 mW), 561 nm (15 mW); HeNe 633 nm (10 mW)). Pictures were acquired with ideal sampling with a PL APO 20x/0.75 or 63x/1.4 CS2 objectives. The images were processed with Icy software.

### Neuronal imaging

All experiments on mice were performed in accordance with European Community guidelines legislation and reviewed by the local ethical committee of the Paris Diderot University. Hippocampal cultures from 18-day-old Sprague-Dawley rat embryos were prepared as described previously^49, 75^. Cells were dissociated by treatment with 0.25% trypsin for 15 min at 37°C and plated on poly-Ornithine (1 mg/mL, Sigma) coated glass coverslips in MEM supplemented with 10% horse serum, 0,6% glucose, 0,2% sodium bicarbonate, 2 mM glutamine, and 10 IU/mL penicillin-streptomycin. After attachment neurons were grown in neurobasal medium supplemented with B27 (ThermoFisher) conditioned on astroglia. Neurons were then image either on Leica DMI8 spinning disc microscope for live cells or on SP5 confocal microscope and DMRE microscope for fixed samples. Neurons were incubated 10 min with MemBright at 37°C, washed 3 times in Krebs Ringer buffer, and then fixed with 4% PFA, 0.1% glutaraldehyde during 10 min at 37°C. Samples were then permeabilized if mentioned with 0.1% triton for 4 minutes and incubated sequentially with primary and secondary antibodies. Video-imaging of hippocampal neurons (first day *in vitro)* was performed on a Zeiss 880 confocal on gasp detectors, with 561 nm laser line set at 0.2%. Image were acquired every 2 min during 13 hours in Krebs ringer medium containing 200 nM MemBright Cy3.5. The signal in supplementary movie 5 was not processed neither for the noise nor the photo-bleaching (raw data).

### dSTORM

HeLa cells were cultured as mentioned above. The cells were fixed with 4% PFA during 10 minutes. The PM labelling was performed by incubating the cells with 20 nM MB-Cy3.5 in Opti-MEM during 10 minutes. All non-bound dyes were removed by 3 consecutive washing with PBS. dSTORM imaging was performed on a home built setup based on a Olympus IX-71 inverted microscope with a high-numerical aperture (NA) TIRF objective (Apo TIRF 100 ×, oil, NA 1.49, Olympus).^76^ The samples were imaged in a photoswitching buffer containing 100 mM MEA and an oxygen scavenging system (0.5 mg/mL glucose oxidase, 40 μg/mL catalase, 10% glucose) in PBS. All chemicals were purchased from Sigma-Aldrich. Samples were illuminated with a 532 nm laser (75 mW, Cobolt) and a 405 nm laser (25 mW, Spectra Physics) was used for the activation of the fluorophores. The signal was recorded on an EM-CCD camera (ImagEM, Hamamatsu) (0.106 μm pixel size) with a typical integration time of 8.7 ms. The acquired stacks were processed with ImageJ software^77^ and the dSTORM images were reconstructed from a series of 6000 frames using the Thunder STORM plug-in.^78^ Acquired images were filtered with a wavelet filter and the approximate fluorophore positions were found by setting the threshold value equal to the standard deviation of the 1st wavelet function. Then, precise subpixel localizations were determined by fitting the spots with the Gaussian PSF by weighted least squares method. High-resolution images were reconstructed from obtained localizations and plotted as a histogram with 20 nm pixel size. Stochastic optical reconstruction microscopy (STORM) neuronal imaging was performed on a Vutara microscope (Bruker) with a high-numerical aperture (NA) objective (60x, water, NA 1.2, Olympus). 170 nm coverslips (Menzel glaser 18 mm diameter #1.5) were mounted on a glass slide with a 15 mm hole. The hole was filled with imaging buffer (Tris 50 mM, NaCl 10 mM, 10% glucose, 100 mM MEA, 70 U/mL glucose oxidase (Sigma G0543), 20 g/mL catalase) and sealed with Picodent twinsil. Samples were illuminated successively with a 647 and 561 nm laser and a 405 nm laser for the reactivation of the fluorophores. Neurons were isolated with wide field mosaic microscopy (Cool snap camera) and then imaged for STORM for a series of 30,000-100,000 images with a FLASH4 CMOS camera (20 ms, 20 × 20 microns). 3D-STORM imaging was performed using the bi-plane module allowing the localisation in the xyz direction. The Srx software (Bruker) was used to localise particles in 3D. Correction for chromatic aberration in super-resolution microscopy has been done using multispectral (blue/green/orange/dark red) Tetraspeck beads (Thermofisher T7279) in Bruker’s software.

## Acknowledgements

This work was supported by the European Research Council ERC Consolidator grant BrightSens 648528 and ANR grant (ANR-10-INBS-04). We thank Doriane Heimburger for the KB cell culture, Romain Vauchelles for his assistance at the PIQ platform, Pauline Meyer for the mass measurements, Bohdan Wasylyk for providing the KB cells and Dr. Florence Toti and Mohamad Kassem for providing fresh rat liver tissues. We acknowledge the ImagoSeine core facility of the Institut Jacques Monod, member of IBiSA and the France-BioImaging infrastructure. We thank Dr. Carl G. Ebeling (Worldwide Application Scientist for Bruker Fluorescence Microscopy) for his help with the use of the Bruker Vutara system and its SRX visualisation and analysis software for rendering localisation data. We thank the NeurImag imaging facility of the Institute of Psychiatry and Neuroscience of Paris, and the national research group in Microscopy of the living (GDR ImaBio) for promoting meetings allowing inter-disciplinary collaborations.

## Author contributions

MC and ASK planned the project. MC synthesized, purified and characterized the MemBright and led the spectroscopy experiments with PA and ASK. PA studied the spectroscopic and two-photon properties. Cell imaging was performed by MC, PA and HA (HeLa d-STORM). Two-photon tissue imaging was realized by OF and LD. EB led the cytotoxicity evaluation. LD led the tissue and neuronal imaging as well as the super resolution imaging. YM, TG and ASK contributed materials and analysis tools. MC, LD and ASK wrote the manuscript with some assistance from other co-authors.

## References

1. Lingwood, D. & Simons, K. Lipid Rafts As a Membrane-Organizing Principle. Science 327, 46–50 (2010).

2. Rello, S. et al. Morphological criteria to distinguish cell death induced by apoptotic and necrotic treatments. Apoptosis 10, 201–208 (2005).

3. Matsuzaki, M., Honkura, N., Ellis-Davies, G. C. R. & Kasai, H. Structural basis of long-term potentiation in single dendritic spines. Nature 429, 761–766 (2004).

4. Lavis, L. D. & Raines, R. T. Bright Ideas for Chemical Biology. ACS Chem. Biol. 3, 142–155 (2008).

5. Fernández-Suárez, M. & Ting, A. Y. Fluorescent probes for super-resolution imaging in living cells. Nat. Rev. Mol. Cell Biol. 9, 929–943 (2008).

6. Neto, B. A. D., Corrêa, J. R. & Silva, R. G. Selective mitochondrial staining with small fluorescent probes: importance, design, synthesis, challenges and trends for new markers. RSC Adv. 3, 5291–5301 (2013).

7. Collot, M. et al. Ultrabright and Fluorogenic Probes for Multicolor Imaging and Tracking of Lipid Droplets in Cells and Tissues. J. Am. Chem. Soc. (2018). doi:10.1021/jacs.7b12817

8. Arai, S., Lee, S.-C., Zhai, D., Suzuki, M. & Chang, Y. T. A Molecular Fluorescent Probe for Targeted Visualization of Temperature at the Endoplasmic Reticulum. Sci. Rep. 4, 6701 (2014).

9. Life Technologies. available at: https://www.thermofisher.com/.

10. Despras, G. et al. H-Rubies, a new family of red emitting fluorescent pH sensors for living cells. Chem. Sci. 6, 5928–5937 (2015).

11. Chen, X. et al. Lysosomal Targeting with Stable and Sensitive Fluorescent Probes (Superior LysoProbes): Applications for Lysosome Labeling and Tracking during Apoptosis. Sci. Rep. 5, 9004 (2015).

12. Klymchenko, A. S. & Kreder, R. Fluorescent Probes for Lipid Rafts: From Model Membranes to Living Cells. Chem. Biol. 21, 97–113 (2014).

13. Owen, D. M., Williamson, D. J., Magenau, A. & Gaus, K. Sub-resolution lipid domains exist in the plasma membrane and regulate protein diffusion and distribution. Nat. Commun. 3, 1256 (2012).

14. Ziomkiewicz, I. et al. Dynamic conformational transitions of the EGF receptor in living mammalian cells determined by FRET and fluorescence lifetime imaging microscopy. Cytometry A 83, 794–805 (2013).

15. Solanko, L. M. et al. Membrane Orientation and Lateral Diffusion of BODIPY-Cholesterol as a Function of Probe Structure. Biophys. J. 105, 2082–2092 (2013).

16. Honigmann, A. et al. Scanning STED-FCS reveals spatiotemporal heterogeneity of lipid interaction in the plasma membrane of living cells. Nat. Commun. 5, 5412 (2014).

17. Niko, Y., Didier, P., Mely, Y., Konishi, G. & Klymchenko, A. S. Bright and photostable push-pull pyrene dye visualizes lipid order variation between plasma and intracellular membranes. Sci. Rep. 6, 18870 (2016).

18. Kreder, R. et al. Blue fluorogenic probes for cell plasma membranes fill the gap in multicolour imaging. Rsc Adv. 5, 22899–22905 (2015).

19. Shynkar, V. V. et al. Fluorescent Biomembrane Probe for Ratiometric Detection of Apoptosis. J. Am. Chem. Soc. 129, 2187–2193 (2007).

20. Kucherak, O. A. et al. Switchable Nile Red-Based Probe for Cholesterol and Lipid Order at the Outer Leaflet of Biomembranes. J. Am. Chem. Soc. 132, 4907–4916 (2010).

21. Kwiatek, J. M. et al. Characterization of a New Series of Fluorescent Probes for Imaging Membrane Order. Plos One 8, e52960 (2013).

22. López-Duarte, I., Vu, T. T., Izquierdo, M. A., Bull, J. A. & Kuimova, M. K. A molecular rotor for measuring viscosity in plasma membranes of live cells. Chem. Commun. 50, 5282–5284 (2014).

23. Zhang, X., Wang, C., Jin, L., Han, Z. & Xiao, Y. Photostable Bipolar Fluorescent Probe for Video Tracking Plasma Membranes Related Cellular Processes. ACS Appl. Mater. Interfaces 6, 12372–12379 (2014).

24. Heek, T. et al. An Amphiphilic Perylene Imido Diester for Selective Cellular Imaging. Bioconjug. Chem. 24, 153–158 (2013).

25. Dal Molin, M. et al. Fluorescent Flippers for Mechanosensitive Membrane Probes. J. Am. Chem. Soc. 137, 568–571 (2015).

26. Fin, A., Jentzsch, A. V., Sakai, N. & Matile, S. Oligothiophene Amphiphiles as Planarizable and Polarizable Fluorescent Membrane Probes. Angew. Chem.-Int. Ed. 51, 12736–12739 (2012).

27. Cui, Q. et al. Binding-Directed Energy Transfer of Conjugated Polymer Materials for Dual-Color Imaging of Cell Membrane. Chem. Mater. 28, 4661–4669 (2016).

28. Wang, H.-Y. et al. Imaging plasma membranes without cellular internalization: multisite membrane anchoring reagents based on glycol chitosan derivatives. J. Mater. Chem. B 3, 6165–6173 (2015).

29. Jiang, Y.-W. et al. In Situ Visualization of Lipid Raft Domains by Fluorescent Glycol Chitosan Derivatives. Langmuir ACS J. Surf. Colloids 32, 6739–6745 (2016).

30. Yuan, L., Lin, W., Zheng, K., He, L. & Huang, W. Far-red to near infrared analyte-responsive fluorescent probes based on organic fluorophore platforms for fluorescence imaging. Chem. Soc. Rev. 42, 622–661 (2012).

31. Umezawa, K., Citterio, D. & Suzuki, K. New Trends in Near-Infrared Fluorophores for Bioimaging. Anal. Sci. 30, 327–349 (2014).

32. Collot, M. et al. Bright fluorogenic squaraines with tuned cell entry for selective imaging of plasma membrane vs. endoplasmic reticulum. Chem. Commun. 51, 17136–17139 (2015).

33. Gonçalves, M. S. T. Fluorescent Labeling of Biomolecules with Organic Probes. Chem. Rev. 109, 190–212 (2009).

34. Hou, T.-C., Wu, Y.-Y., Chiang, P.-Y. & Tan, K.-T. Near-infrared fluorescence activation probes based on disassembly-induced emission cyanine dye. Chem. Sci. 6, 4643–4649 (2015).

35. van de Linde, S. et al. Direct stochastic optical reconstruction microscopy with standard fluorescent probes. Nat. Protoc. 6, 991–1009 (2011).

36. Würthner, F., Kaiser, T. E. & Saha-Möller, C. R. J-Aggregates: From Serendipitous Discovery to Supramolecular Engineering of Functional Dye Materials. Angew. Chem. Int. Ed. 50, 3376–3410 (2011).

37. Gousset, K. et al. Prions hijack tunnelling nanotubes for intercellular spread. Nat. Cell Biol. 11, 328–336 (2009).

38. Helmchen, F. & Denk, W. Deep tissue two-photon microscopy. Nat. Methods 2, 932–940 (2005).

39. Svoboda, K. & Yasuda, R. Principles of Two-Photon Excitation Microscopy and Its Applications to Neuroscience. Neuron 50, 823–839 (2006).

40. Yao, S. & Belfield, K. D. Two-Photon Fluorescent Probes for Bioimaging. Eur. J. Org. Chem. 2012, 3199–3217 (2012).

41. Lukomska, J. et al. Two-photon induced fluorescence of Cy5-DNA in buffer solution and on silver island films. Biochem. Biophys. Res. Commun. 328, 78–84 (2005).

42. Kobat, D. et al. Deep tissue multiphoton microscopy using longer wavelength excitation. Opt. Express 17, 13354 (2009).

43. Albota, M. A., Xu, C. & Webb, W. W. Two-photon fluorescence excitation cross sections of biomolecular probes from 690 to 960 nm. Appl. Opt. 37, 7352 (1998).

44. Kim, H. M. & Cho, B. R. Small-Molecule Two-Photon Probes for Bioimaging Applications. Chem. Rev. 115, 5014–5055 (2015).

45. Guo, L. et al. Styrylpyridine salts-based red emissive two-photon turn-on probe for imaging the plasma membrane in living cells and tissues. Analyst 141, 3228–3232 (2016).

46. Rasia-Filho, A. R.-F. Dendritic spines observed by extracellular Dil dye and immunolabeling under confocal microscopy. Protoc. Exch. (2010). doi:10.1038/nprot.2010.153

47. Feng, G. et al. Imaging Neuronal Subsets in Transgenic Mice Expressing Multiple Spectral Variants of GFP. Neuron 28, 41–51 (2000).

48. Ricard, C. & Debarbieux, F. C. Six-color intravital two-photon imaging of brain tumors and their dynamic microenvironment. Front. Cell. Neurosci. 8, 57 (2014).

49. Danglot, L., Triller, A. & Bessis, A. Association of gephyrin with synaptic and extrasynaptic GABAa receptors varies during development in cultured hippocampal neurons. Mol. Cell. Neurosci. 23, 264–278 (2003).

50. Revelo, N. H. et al. A new probe for super-resolution imaging of membranes elucidates trafficking pathways. J Cell Biol 205, 591–606 (2014).

51. Revelo, N. H. & Rizzoli, S. O. The Membrane Marker mCLING Reveals the Molecular Composition of Trafficking Organelles. in Current Protocols in Neuroscience (John Wiley & Sons, Inc, 2001). doi:10.1002/0471142301.ns0225s74

52. Dequidt, C. et al. Fast Turnover of L1 Adhesions in Neuronal Growth Cones Involving Both Surface Diffusion and Exo/Endocytosis of L1 Molecules. Mol. Biol. Cell 18, 3131–3143 (2007).

53. Kasai, H., Fukuda, M., Watanabe, S., Hayashi-Takagi, A. & Noguchi, J. Structural dynamics of dendritic spines in memory and cognition. Trends Neurosci. 33, 121–129 (2010).

54. Huang, B., Babcock, H. & Zhuang, X. Breaking the Diffraction Barrier: Super-Resolution Imaging of Cells. Cell 143, 1047–1058 (2010).

55. Rust, M. J., Bates, M. & Zhuang, X. Sub-diffraction-limit imaging by stochastic optical reconstruction microscopy (STORM). Nat. Methods 3, 793–795 (2006).

56. Betzig, E. et al. Imaging Intracellular Fluorescent Proteins at Nanometer Resolution. Science 313, 1642–1645 (2006).

57. Hess, S. T., Girirajan, T. P. K. & Mason, M. D. Ultra-high resolution imaging by fluorescence photoactivation localization microscopy. Biophys. J. 91, 4258–4272 (2006).

58. Porrero, C., Rubio-Garrido, P., Avendaño, C. & Clascá, F. Mapping of fluorescent protein-expressing neurons and axon pathways in adult and developing Thy1-eYFP-H transgenic mice. Brain Res. 1345, 59–72 (2010).

59. Horikawa, K. & Armstrong, W. E. A versatile means of intracellular labeling: injection of biocytin and its detection with avidin conjugates. J. Neurosci. Methods 25, 1–11 (1988).

60. Honig, M. G. & Hume, R. I. Fluorescent carbocyanine dyes allow living neurons of identified origin to be studied in long-term cultures. J. Cell Biol. 103, 171–187 (1986).

61. Lanciego, J. L. & Wouterlood, F. G. A half century of experimental neuroanatomical tracing. J. Chem. Neuroanat. 42, 157–183 (2011).

62. Chevalier, A. C. & Rosenberger, T. A. Increasing acetyl-CoA metabolism attenuates injury and alters spinal cord lipid content in mice subjected to experimental autoimmune encephalomyelitis. J. Neurochem. 141, 721–737 (2017).

63. Whitney, M. A. et al. Fluorescent peptides highlight peripheral nerves during surgery in mice. Nat. Biotechnol. 29, 352–356 (2011).

64. Dani, A., Huang, B., Bergan, J., Dulac, C. & Zhuang, X. Superresolution Imaging of Chemical Synapses in the Brain. Neuron 68, 843–856 (2010).

65. Herrmannsdörfer, F. et al. 3D d STORM Imaging of Fixed Brain Tissue. Methods Mol. Biol. Clifton NJ 1538, 169–184 (2017).

66. Beghin, A. et al. Localization-based super-resolution imaging meets high-content screening. Nat. Methods 14, 1184–1190 (2017).

67. Lagache, T. et al. Mapping molecular assemblies with fluorescence microscopy and object-based spatial statistics. Nat. Commun. 9, 698 (2018).

68. Levet, F. et al. SR-Tesseler: a method to segment and quantify localization-based super-resolution microscopy data. Nat. Methods 12, 1065–1071 (2015).

69. Hope, M. J., Bally, M. B., Webb, G. & Cullis, P. R. Production of large unilamellar vesicles by a rapid extrusion procedure: characterization of size distribution, trapped volume and ability to maintain a membrane potential. Biochim. Biophys. Acta 812, 55–65 (1985).

70. Rurack, K. & Spieles, M. Fluorescence Quantum Yields of a Series of Red and Near-Infrared Dyes Emitting at 600-1000 nm. Anal. Chem. 83, 1232–1242 (2011).

71. Texier, I. et al. Cyanine-loaded lipid nanoparticles for improved in vivo fluorescence imaging. J. Biomed. Opt. 14, 054005–054005-11 (2009).

72. Alessi, A., Salvalaggio, M. & Ruzzon, G. Rhodamine 800 as reference substance for fluorescence quantum yield measurements in deep red emission range. J. Lumin. 134, 385–389 (2013).

73. Benson, R. & Kues, H. Fluorescence Properties of Indocyanine Green as Related to Angiography. Phys. Med. Biol. 23, 159–163 (1978).

74. Clamme, J. P., Azoulay, J. & Mély, Y. Monitoring of the Formation and Dissociation of Polyethylenimine/DNA Complexes by Two Photon Fluorescence Correlation Spectroscopy. Biophys. J. 84, 1960–1968 (2003).

75. Banker, G. A. & Cowan, W. M. Rat hippocampal neurons in dispersed cell culture. Brain Res. 126, 397–342 (1977).

76. Kempf, N., Didier, P., Postupalenko, V., Bucher, B. & Mély, Y. Internalization mechanism of neuropeptide Y bound to its Y 1 receptor investigated by high resolution microscopy. Methods Appl. Fluoresc. 3, 025004 (2015).

77. Schneider, C. A., Rasband, W. S. & Eliceiri, K. W. NIH Image to ImageJ: 25 years of image analysis. Nat. Methods 9, 671–675 (2012).

78. Ovesný, M., Křížek, P., Borkovec, J., Švindrych, Z. & Hagen, G. M. ThunderSTORM: a comprehensive ImageJ plug-in for PALM and STORM data analysis and super-resolution imaging. Bioinformatics 30, 2389–2390 (2014).

